# CRISPRi-mediated validation of candidate *Mycobacterium abscessus* drug targets during host infection

**DOI:** 10.64898/2025.12.16.694790

**Authors:** Rashmi Gupta, Breven S. Simcox, Kyle H. Rohde

## Abstract

*Mycobacterium abscessus* (*Mab*) is a multidrug-resistant nontuberculous mycobacterium that causes debilitating TB-like pulmonary infections for which effective treatment options are lacking. Poor *in vivo* drug efficacy may stem from altered vulnerability of drug targets driven by host-specific environmental conditions. To enable validation and prioritization of candidate drug targets *in vivo*, we exploited CRISPRi (CRi) gene silencing in multiple mouse infection models. Inducible silencing of *ftsZ*_*Mab*_, a previously validated target, and three predicted targets (*leuS*_*Mab*_, *folP*_*Mab*_, *fusA*_*Mab*_) confirmed their essentiality *in vitro*. We then assessed the *in vivo* vulnerability of these targets in both immunocompetent C57BL/6N and immunodeficient NSG mice by assessing the impact of CRi silencing on pulmonary mycobacterial burden. In NSG mice, silencing of all four genes led to comparable decreases in *Mab* burden. However, in C57BL/6N mice, the degree of *Mab* clearance varied among targets, suggesting that immune pressure may influence the outcome of CRi-mediated gene silencing. Notably, repression of *fusA*_*Mab*_ yielded a larger decline in mycobacterial burden in C57BL/6N mice despite a lower level of gene silencing *in vitro*, consistent with enhanced vulnerability of this target. Overall, this study demonstrated that *ftsZ*_*Mab*_, *leuS*_*Mab*_, *folP*_*Mab*_, and *fusA*_*Mab*_ are essential for *Mab* growth *in vitro* and, for the first time, validated their vulnerability to inhibition by CRi during infection. These data also identified potential context-dependent target vulnerabilities, which could inform the prioritization of bacterial drug targets and accelerate the development of effective therapeutics for *Mab* infections.

## INTRODUCTION

The rising incidence of antibiotic resistance among bacterial pathogens poses a grave threat to global public health, necessitating the development of innovative therapeutic strategies. *Mycobacterium abscessus* (*Mab*), which causes chronic pulmonary infections, particularly in individuals with cystic fibrosis, COPD, bronchiectasis and other co-morbidities (1-7), has emerged as an almost untreatable opportunistic pathogen. This is due to its intrinsic resistance to multiple antibiotics, including the ones used against TB infections, resulting from a highly impermeable cell wall, efficient efflux pump systems, and inducible resistance mechanisms which confer resistance to a broad spectrum of antibiotics (8-11). *Mab* infections require a prolonged, aggressive treatment regimen with multiple oral and injectable antibiotics with failure rates that remain unacceptably high (up to 70%) (12, 13). In addition to the paucity of antibiotics able to effectively kill *Mab* even *in vitro*, discrepancies between *in vitro* and *in vivo* drug efficacy attributable to host-induced physiological changes in *Mab* pose an additional challenge. This highlights the need to evaluate the vulnerability of candidate drug targets within the context of infection. In recent years, CRISPR interference (CRi) has emerged as a powerful tool for gene manipulation, offering precise control over gene expression through transcriptional silencing of targeted genes. By utilizing a catalytically inactive variant of the CRISPR-associated protein Cas9 (dCas9) coupled with sequence-specific guide RNAs (sgRNAs), CRi enables efficient and reversible silencing of target genes upon induction with anhydrotetracycline (ATc) without inducing permanent genetic changes (14, 15). Several research groups including ours have utilized this tool in *Mab* to probe gene functions including those essential for bacterial growth and drug resistance (16-20). CRi has numerous applications from studying genotype-phenotype relationships, targeting multiple genes to identify functional synthetic lethality, engineering metabolic pathways to optimize synergistic interactions and identifying disease mechanisms (21-28). Applied to *M. tuberculosis*, CRi technology revealed the differential vulnerability of essential genes to silencing *in vitro*, which could serve as a proxy indicator of the vulnerability or “druggability” of drug targets (29). Evaluation of gene silencing *in vivo* will help us understand physiological relevance and likely effectiveness of inhibitors of specific targets in a clinical context.

The combination of technologies like CRi with animal models provides a valuable strategy for assessing the impact of *in vivo* gene silencing on bacterial virulence, host-pathogen interactions, and potential therapeutic interventions. Several studies have focused on mycobacterial gene expression modulation in animals, primarily utilizing TetR-based systems which operate independently of CRi-mediated mechanism for regulating *Mtb* genes (30-37). However, the role of *Mab* genes including virulence factors and potential drug targets *in vivo* remains a significant knowledge gap due to delayed implementation of genetic tools in *Mab* and lack of animal modes of persistent *Mab* infection (38, 39). To date, only a handful of studies involving silencing of *Mab* genes during infection have been reported. One study employed the Tet-OFF regulatory system to modulate expression of the *mmpL3* mycolic acid transporter gene to confirm its essentiality in *Mab* during zebrafish infection (40). This animal model is evolutionarily distant from humans and thus lacks direct translational relevance to pathogenesis and infection. During the preparation of this manuscript, the reported application of CRi in more relevant murine models (41, 42) highlighted the potential of this approach to yield insights into the therapeutic potential of genes under physiological conditions. The ability to regulate gene expression during an infection would also provide much needed molecular tools to understand *Mab* pathogenesis and virulence.

Previously, we have utilized the mycobacterial CRi platform to validate essential genes and drug-resistant genes in *Mab* (17). In this study, we demonstrated CRi-mediated silencing of *ftsZ*, a previously validated target, during an animal infection using immunodeficient and immunocompetent mouse models. In addition, we validated three additional essential genes (*leuS*_*Mab*_, *fusA*_*Mab*_, *folP*_*Mab*_) and evaluated their vulnerability to lethal silencing both *in vitro* and *in vivo*. Defining the differential vulnerability of essential genes, which represent candidate drug targets, to genetic silencing *in vivo* would be valuable towards validating and prioritizing new drug candidates for therapeutics development.

## RESULTS & DISCUSSION

### CRi -mediated validation of predicted drug targets

In our previous study, we utilized the mycobacterial single plasmid pLJR962 CRi platform optimized for *M. smegmatis* featuring dCas9_Sth1_ (43) and demonstrated silencing and essentiality of *ftsZ*_*Mab*_ (17). Here, we extended our studies by evaluating CRi silencing of three additional genes with crucial roles in protein and nucleotide synthesis - *fusA*_*Mab*_ encoding elongation factor G (EF-G) (MAB_3849c), *leuS*_*Mab*_ coding for leucine tRNA synthetase (MAB_4923c), and *folP*_*Mab*_, a folate dihydropteroate synthase (MAB_0535). These targets are predicted to be essential based on Tn-seq analyses and corroborated by homology with essential *Mtb* genes as well as the antimicrobial efficacy of target-specific inhibitors against *Mtb* (44-49). We generated CRi constructs designed to hybridize adjacent to protospacer adjacent motifs (PAMs) with the highest predicted strength as described previously (**Table S1**) (17). After introduction into *Mab*, the impact of silencing on mRNA levels of the targeted gene and subsequent loss in cell viability (**Fig. 1)** were evaluated. Induction with either a low (200ng/ml) or high dose (10µg/ml) of ATc resulted in dose-dependent reduction in mRNA levels for all target genes except for *fusA*_*Mab*_ (**Fig. 1A**). At the low ATc dose, transcript levels for three of the target genes (*ftsZ*_*Mab*_, *leuS*_*Mab*_, *folP*_*Mab*_) were reduced ∼10-fold, whereas *fusA*_*Mab*_ silencing was less effective, with only ∼4-fold decrease. The higher dose of ATc yielded ∼100-fold reduction in *ftsZ*_*Mab*_, *leuS*_*Mab*_, and *folP*_*Mab*_ transcripts, but no additional decline in *fusA*_*Mab*_ mRNA levels was noted. As expected, the reduction in transcript levels corresponded with cell viability loss following CRi induction. All the targets including *fusA*_*Mab*_ exhibited a ∼3-4 log decrease in CFU (**Fig. 1B, Fig. S1**). The same level of bactericidal activity yielded by CRi targeting *fusA*_*Mab*_ despite much weaker effect on mRNA levels would suggest *fusA*_*Mab*_ is a more vulnerable target. Overall, the strong correlation between transcriptional silencing and decreased *Mab* viability validates *leuS*_*Mab*_, *folP*_*Mab*_, and *fusA*_*Mab*_ as essential genes and promising drug targets. However, their individual vulnerabilities may vary and become more pronounced under disease relevant conditions, making it crucial to identify and quantify these differences important for drug target prioritization.

**Figure 1:**
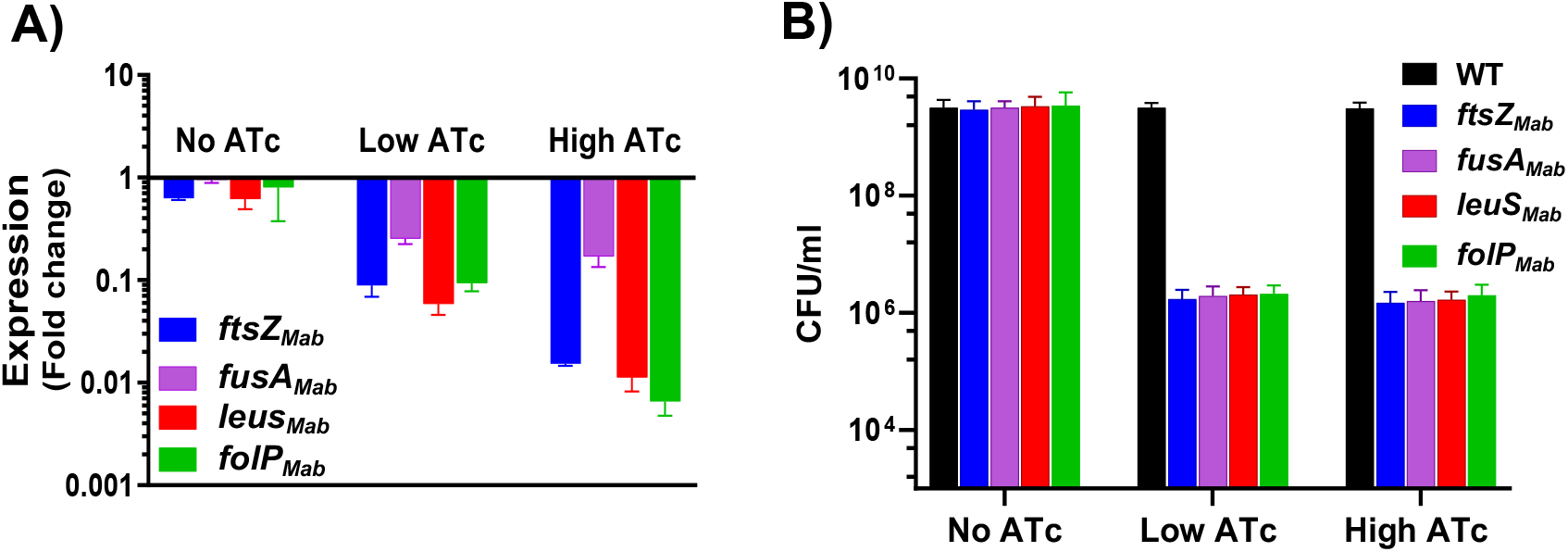
Impact of CRi silencing on predicted *Mab* essential genes. **(A)** mRNA levels relative to uninduced WT control. **(B)** CFU/ml upon ATc induction at low (200ng/ml) and high (10µg/ml) ATc concentration. N=3

### Optimizing intranasal infection route of *Mab* mouse model

Assessment of gene vulnerabilities based on CRi silencing *in vivo* requires a murine model for *Mab* infection. To establish *Mab* pulmonary infection in mouse lungs, the intranasal delivery method was adopted as it is simple and mimics the natural infection route. However, pilot mouse infection studies administering *Mab* via intranasal route proved to be variable, including some mice failing to develop an infection indicating suboptimal infection efficiency. Such inconsistency of lung colonization could be interpreted erroneously as bacterial clearance. Several variables including animal body position, and dosing volume could influence reproducibility of intranasal infection (50-52), with higher intranasal instillation volumes reported to yield significant improvements in the infection efficiency of *Fransciella tularensis* (50). Therefore, to establish optimal infection conditions prior to attempting CRi gene silencing *in vivo*, we evaluated the effect of inoculum volume and animal positioning on *Mab* infection efficiency which has not been previously described for *Mab* infection **(Fig. 2A)**. *Mab* was administered intranasally in volumes of 20 and 40 µl in C57BL/6N mice (8–12 weeks old). For each dosing volume, we also assessed body positioning by infecting animals in both supine (S) and vertical (V) positions. The position of the mice during intranasal delivery of *Mab* did not affect infection efficiency, regardless of the inoculum volume (20 or 40 µl) (**Fig. 2B**). In contrast to body positioning, a significant difference (p< 0.005) was noted in infection efficiency between mice dosed with a volume of 20 µl and those with 40 µl 24 h post infection (**Fig. 2B**). Specifically, 100% of 18 mice intranasally administered a 40 µl dose were successfully infected as compared to 78% infection rate in those given a 20 µl dose. Increasing the volume likely increased the probability of the inoculum reaching lungs as opposed to being retained in the upper respiratory airways (51). These findings demonstrate that intranasal instillation volume has a significant impact on the efficiency of pulmonary delivery of *Mab*. Based on the results, subsequent experiments in this study were conducted using *Mab* inoculum volume of 40 µl.

**Figure 2:**
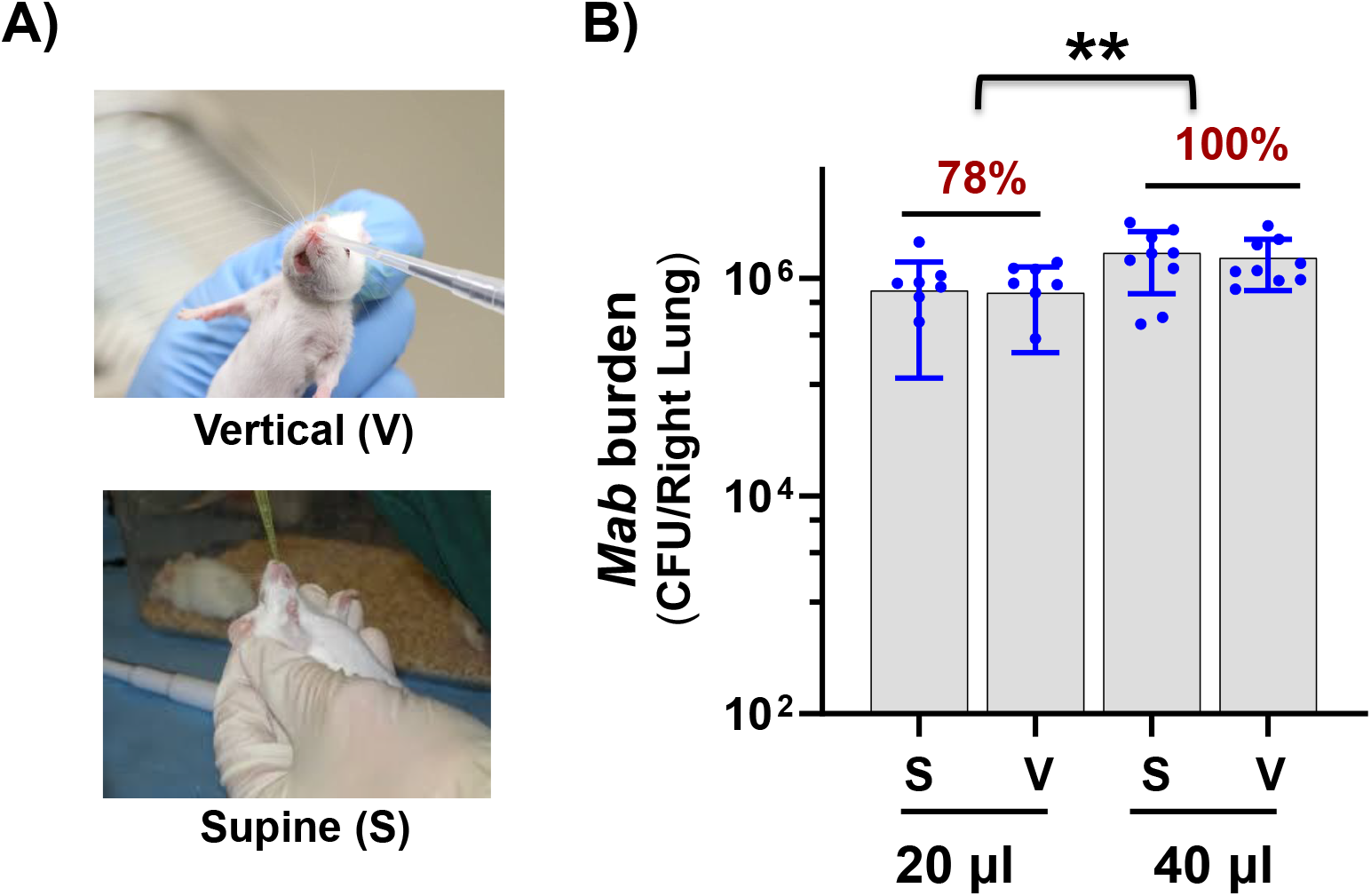
Impact of dosing volume and body position on *Mab* infection efficiency. **(A)** A schematic showing two body positions-vertical (V) and supine (S) evaluated during intranasal infection. It is to be noted that animals used are black-coat C57BL/6N mice and not white coat-colored animals as depicted. **(B)** *Mab* load in the lungs of infected animals when infected with 10^6^ CFU at indicated dosing volumes when held in supine or vertical position. The data is from a total of 9 animals per group from 3 pooled experiments. ** P=0.001, unpaired t-tests with Welch’s correction.

### *In vivo* CRi-mediated silencing in *Mab*

We then applied this optimized model of pulmonary *Mab* infection towards two goals: 1) demonstrating the feasibility of CRi-mediated gene silencing *in vivo* with simple ATc induction via drinking water, and 2) Assessing whether immune pressure influences the vulnerability of different targets by comparing outcomes in immunocompetent C57BL/6N and immunodeficient NSG mice. The presence or absence of an intact immune response in these models may alter the growth and physiology of *Mab* and/or impose stresses that affect the impact of silencing certain genes.

For proof of principle studies to test the feasibility of CRi-mediated gene silencing *in vivo*, we selected a previously validated essential target, *ftsZ*_*Mab*_ (17). NSG (NOD SCID gamma) mice, which lack mature T cells, B cells, and functional NK (natural killer) cells, represent a simple model to study the effect of CRi gene silencing on *Mab* pulmonary burden without clearance mediated by innate and adaptive immunity. Immunodeficient NSG mice were infected with the *ftsZ*_*Mab*_ CRi strain (or empty CRi vector) using the infection scheme outlined in **Fig. 3A**. Twenty-four hours post-infection, both the control and the *ftsZ* CRi groups of mice were provided drinking water supplemented with ATc inducer (100 and 200 µg/ml) and sucrose (to mask the taste of ATc and facilitate sufficient intake of ATc-laced water). Doxycycline is the more commonly used TetR inducer in mouse models due to its established pharmacokinetics and tissue distribution (32, 53, 54). However, given that recent studies have shown that ATc does mediate robust TetR-dependent gene repression in mice (41, 42, 54), we used ATc to maintain consistency with *in vitro* studies while avoiding concerns about the antimicrobial effects associated with doxycycline. When induced with 100 µg/ml ATc, the control group maintained a stable bacterial burden whereas CRi silencing of *ftsZ*_*Mab*_ resulted in a 0.7 log decline relative to the control on day 3, compared to a 1-log decrease in CFU when mice were administered 200 µg/ml ATc (**Fig. 3B)**. The higher ATc dose was chosen for further studies based on the higher CFU decline noted although the difference between the two doses was not statistically significant. Thereafter, we infected NSG mice with other CRi strains (*leuS*_*Mab*_, *folP*_*Mab*_ and *fusA*_*Mab*_) and induced with 200 µg/ml ATc. Suppression of these three targets also resulted in a similar CFU decline (∼0.5 log_10_), after 3 days of CRi induction by ATc (**Fig. 3C)**. This short duration allowed ample time observe a decline in *Mab* CFU for rapid assessment of target gene vulnerability while also minimizing the emergence of CRi escape mutants which we noted in previous *in vitro* studies (17).

**Figure 3:**
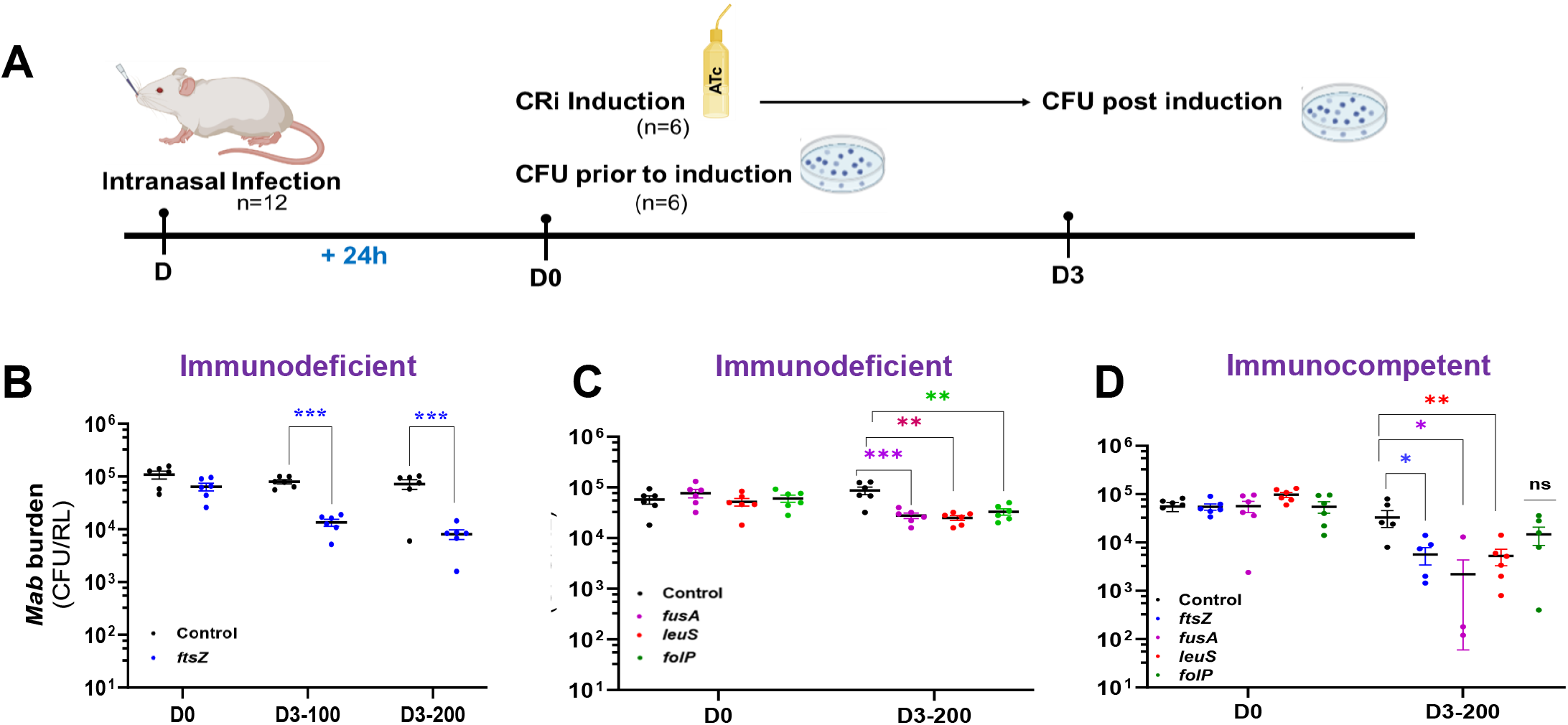
CRISPRi-mediated silencing of *Mab* genes in mice. **(A)** Illustration of the infection scheme. Mice were infected intranasally with control, *ftsZ*_*Mab*_ and other CRi strains (*fusA*_*Mab*_, *leuS*_*Mab*_, and *folP*_*Mab*_) and administered ATc in drinking water for 3 days. Prior to ATc induction at D0 (24h p.i.) and at D3 (3d post-induction) mycobacterial burden in the right lung was determined. NSG mice induced with **(B)** 100 and 200 µg/ml ATc water and **(C)** with 200 µg/ml ATc water. **(D)** C57BL/6N mice induced with 200 µg/ml ATc water. Data are mean ±SEM of 6 mice per group per time point. Statistical analysis: (B) two-way ANOVA with Tukey’s multiple comparisons, (C and D)-One-way ANOVA with Dunnett’s comparisons, * P≤0.01, ** P≤0.001, ** *P≤0.0001.

Next, we sought to determine the impact of CRi silencing on *Mab* growth and survival during infection in immunocompetent mice to assess whether immune pressure or host-derived stresses could influence target vulnerability. In C57BL/6N mice, a ∼0.8-1.0 log_10_ reduction in CFU was noted upon induction of silencing of 3 *Mab* gene targets (*ftsZ*_*Mab*_, *leuS*_*Mab*_, and *fusA*_*Mab*_), whereas the *folP*_*Mab*_ CRi strain showed no statistical decline (0.3 log_10_) in *Mab* burden compared to the control post-induction (D3) (**Fig. 3D)**. *fusA*_*Mab*_ appeared to be notably vulnerable with no *Mab* colonies recovered from 50% of the animals and among those with detectable bacilli, ⅔ mice had ∼2-log lower CFU compared to the single mouse with high burden (∼10^4^ CFU). This highlights higher vulnerability of the *fusA*_*Mab*_ target in context of immune pressure, consistent with our *in vitro* studies. This is also in agreement with *Mtb* vulnerability index based on Tn-seq essentiality and repression outcome predictability (44, 55). The use of immune-compromised and -competent mouse models revealed vulnerability differences in the selected essential targets with *fusA*_*Mab*_ as the most and *folP*_*Mab*_ as the least conditionally susceptible. In summary, the results demonstrate that *ftsZ*_*Mab*_, *leuS*_*Mab*_, *folP*_*Mab*_ and *fusA*_*Mab*_ genes can be silenced during host infection and are essential for growth and thus represent potential drug targets.

## CONCLUSION

*Mab* is a highly drug-resistant human pathogen, and existing treatments have a high failure rate, underscoring the need to identify new, effective treatment options. The identification and validation of new drug targets is one critical strategy to address this need. While high-throughput *in vitro* mycobacterial screens, such as a genome-wide gene expression (55), and Tn-seq analysis (56) have identified essential targets and highlighted species-specific differences in gene essentiality, *in vivo* validation of target vulnerability remains crucial to determine physiological relevance and “druggability” in a clinical context. CRi provides a versatile genetic tool for assessing the relative vulnerability of putative drug targets to inhibition that facilitates prioritization of potential drug targets. Here, we extended our previous *in vitro* study (17) to silence predicted essential *Mab* genes *in vivo* to probe target vulnerabilities in the context of a pulmonary infection. The major findings of this study are 1) validation of the essential function of three additional *Mab* genes-*leuS*_*Mab*_, *folP*_*Mab*_ and *fusA*_*Mab*_, 2) demonstration of CRi-mediated modulation of *Mab* gene expression in mice and 3) evidence for differences in gene vulnerabilities under *in vitro* versus *in vivo* conditions.

This study highlighted the feasibility of CRi-mediated *Mab* genes suppression *in vivo* in two distinct murine models. The expression of selected gene targets was silenced by CRi induction after establishment of *Mab* infection to truly reflect gene silencing within the host environment. This is in contrast to a strategy implemented in recent studies in which target genes were preemptively silenced by ATc induction 3 days before infection (41, 42). This could have confounded assessment of *in vivo* target vulnerability by depleting essential enzymes prior to infection. Comparison of the four essential targets investigated in this study indicated that *fusA*_*Mab*_ was the most vulnerable to silencing both *in vitro* and *in vivo*. Despite much weaker CRi-mediated transcriptional repression of *fusA*_*Mab*_, the loss of viability *in vitro* was comparable to other targets. The target-specific vulnerability differences noted between immunocompetent and immunocompromised murine models suggest that immune pressure or other conditions within the host niche may alter the impact of target inhibition on mycobacterial viability.

Whereas CRi silencing of *fusA*_*Mab*_ in the context of an innate immune response caused a more dramatic reduction in CFU compared to *ftsZ*_*Mab*_ or *leuS*_*Mab*_, silencing of *folP*_*Mab*_ exhibited a dampened effect on *Mab* lung burden in the immunocompetent mice. The discrepant outcomes of CRi gene silencing versus CFU loss *in vitro* or in NSG and C57BL/6N mice emphasize the importance of assessing drug target vulnerability in relevant mammalian *in vivo* models. Application of this type of data to drug target selection would support prioritization of *fusA*_*Mab*_ over *folP*_*Mab*_ based on the outsized effect on *Mab* viability compared to the level of gene silencing. While this approach offers a rapid strategy to triage or rank drug target vulnerability *in vivo*, the short infection duration (3d) is not ideal to study targets associated with *Mab* persistence and adaptation. Efforts to extend the infection period is hampered by the current lack of suitable immunocompetent model of persistent infection, and possible emergence of CRi escape mutants observed in our previous study (17). Ongoing work in the lab is dedicated to addressing these limitations.

In conclusion, the study demonstrated CRi-mediated *Mab* gene silencing during host infection, highlighting the potential of underexplored essential genes as valuable drug targets. CRi tools can serve a significant pathway for identifying truly sensitive (physiologically relevant) targets and facilitating target prioritization. By uncovering novel targets, this approach may ultimately lead to improved therapeutics that address the challenge of drug resistance.

## MATERIALS AND METHODS

### Bacterial strains and culture conditions

Bacterial strains used in this study (**Table 1**) were cultured in Middlebrook 7H9 supplemented with 0.05% Tween 80 and 10% oleic acid/albumin/dextrose/catalase (OADC) and incubated at 37ºC and 5% CO_2_. Kanamycin 50 µg/mL (KAN_50_), Apramycin 50 µg/mL (APRA_50_), cycloheximide 100 µg/mL and amikacin 32 µg/mL (AMK_32_) were added when appropriate, depending on expected resistance profile of the strain. Stock solution of anhydrotetracycline (ATc) was prepared according to the manufacturer’s instructions. Stocks were stored at -20ºC.

**Table 1:**
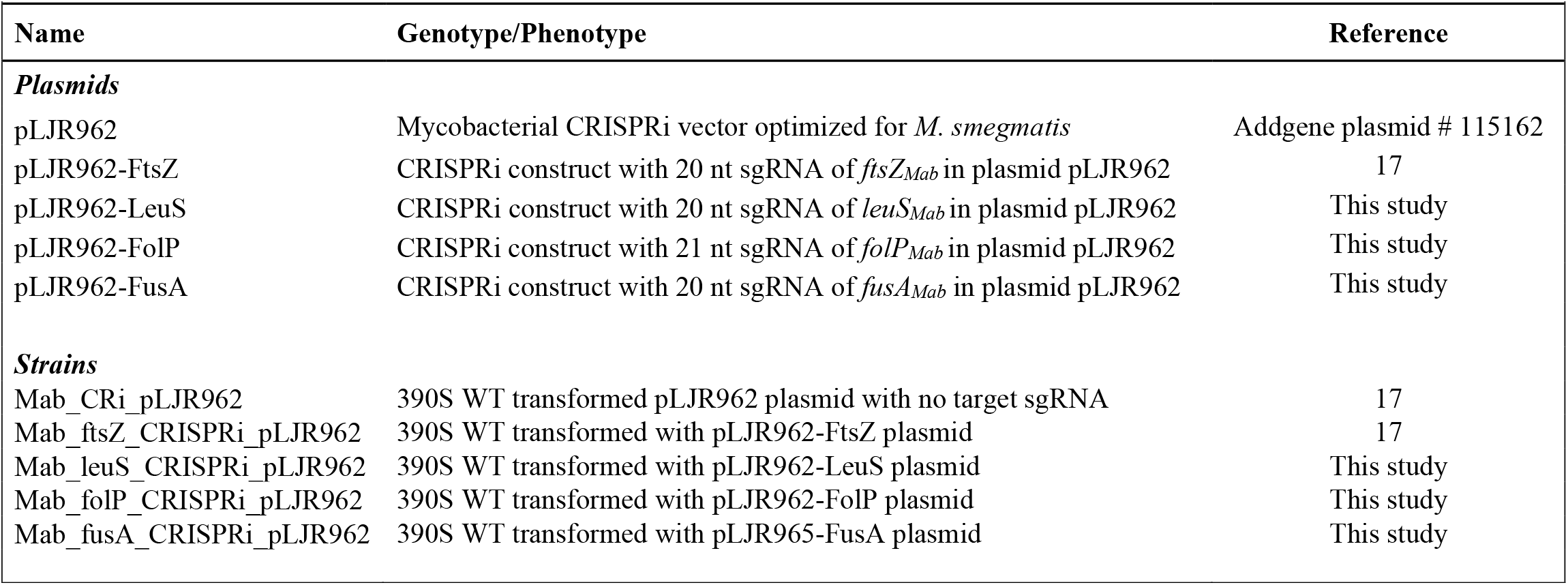
Bacterial Plasmids and Strains.

### Construction of *Mab* CRi knockdowns

We constructed CRi knockdowns in four genes (MAB_0535: dihydropterate synthase, *folP*_Mab_; MAB_4923c: leucine tRNA ligase, *leuS*_Mab_; MAB_2009: Cell division protein, *ftsZ*; and MAB_3849c: Elongation factor G, *fusA*) in pLJR962 (gift from Sarah Fortune, Addgene plasmids #115162 and #11563http://n2t.net/addgene:115162, http://n2t.net/addgene:115163) (43). PAM and sgRNA target sequence selection was carried out as per Rock *et al*. study (43). CRi constructs of *leuS*_Mab_, *folP*_Mab_, *ftsZ*_Mab_, and *fusA*_*Mab*_ were created using a PCR-based mutagenesis strategy as described earlier (17). See **Table S2** for sequences of primers used for CRi plasmid generation, colony screening and sequencing.

### RNA isolation and qRT-PCR

*Mab* CRi strains were grown and induced by ATc for 20h as described before (17). An uninduced sample was kept as the control. RNA isolation was carried out using RNeasy kit as described previously (17, 57). Ct values were normalized against the housekeeping gene, *sigA* and fold change was calculated against an uninduced sample run in parallel using the -2^ΔΔCt^ method. Fold change is represented relative to WT un-induced (no ATc). All qRT-PCR primers are listed in **Table S2**.

### CFU Enumeration

A dilution-plating assay was conducted to determine the effect of CRi mediated repression of the putative essential genes, *ftsZ*_*Mab*_, *leuS*_*Mab*_, *folP*_*Mab*_ and *fusA*_*Mab*_ on cell viability. For the assay, the cultures were grown from freezer stock to log phase (OD_600 =_0.4-0.8), then diluted back to OD_600 =_0.3 and 5 µl of 2-fold serial dilutions were spotted onto 7H10 agar plates with and without ATc. The plates were incubated at 37°C for 5 days and CFU/ml was calculated.

### In vivo CRi silencing

We utilized immunocompetent C57BL6/N and immunodeficient NSG (NOD scid gamma mouse) murine models to assess *Mab* gene silencing during murine infection. C57BL6/N (Taconic Biosciences) and NSG (NOD SCID gamma) mice (UCF breeding colony) were housed in an AAALAC credited UCF Lake Nona vivarium under stringent sterile conditions. Eight to 10-week-old male and female mice were anesthetized with inhaled isoflurane (2.5 L/min), and intranasally infected with an inoculum of 10^5^ CFU of 5 CRi-*Mab* strains, including the control (empty vector) strain. A log-phase culture was syringed 5x using a 27-G needle syringe to disrupt clumps before dilution in 40 µl PBS. Each group consisted of 6 mice per time point with an equal sex distribution (3 males, 3 females). After 24 h of infection, animals were euthanized to assess baseline infection levels which served as D0 controls. Thereafter, mice, including those which received the empty vector were treated with ATc (at indicated concentration) and sucrose (5%) supplemented drinking water in red-tinted water bottles (light protection). After 3 days of ATc induction, lungs were harvested to determine mycobacterial burden. Right lungs were excised and homogenized in 1ml PBS with 1 mm diameter silicon carbide beads (0.5ml) in a beat beater (Biospec) for 2 mins (max. speed). The homogenized lung tissue was then serially diluted and plated (50 µl) onto 7H10 agar plates. After 5-7 days of incubation, colonies were counted, and CFU/ml was calculated.

## Statistical Analysis

A statistical difference was determined using t-tests with pairwise comparisons and ANOVA analyses with appropriate Tukey or Dunnett’s post-test analysis. GraphPad Prism was used for this analysis. Each data point indicates a single animal unless otherwise stated. Statistical tests are indicated in the legends wherever appropriate.

## Author Contribution

**Rashmi Gupta:** Conceptualization; Data curation; Formal analysis; Investigation; Methodology; Visualization; Roles/Writing - original draft; Writing - review & editing. **Breven Gaines:** Methodology. **Kyle H. Rohde:** Conceptualization; Formal analysis; Funding acquisition; Methodology; Project administration; Resources; Supervision; Writing - review & editing.

## Acknowledgements

We are thankful to Maria Tori Buencamino, Sahiba Ahmed, and Alexandria Iakovidis for constructing CRi knockdown plasmids. The study was funded by NIH award R21AI156221 to KHR.

## References

1. Benwill JL, Wallace RJ, Jr. 2014. Mycobacterium abscessus: challenges in diagnosis and treatment. Curr Opin Infect Dis 27:506–10.

2. Brown-Elliott BA, Griffith DE, Wallace RJ, Jr. 2002. Diagnosis of nontuberculous mycobacterial infections. Clin Lab Med 22:911–25, vi.

3. Griffith DE, Aksamit T, Brown-Elliott BA, Catanzaro A, Daley C, Gordin F, Holland SM, Horsburgh R, Huitt G, Iademarco MF, Iseman M, Olivier K, Ruoss S, von Reyn CF, Wallace RJ, Jr., Winthrop K. 2007. An official ATS/IDSA statement: diagnosis, treatment, and prevention of nontuberculous mycobacterial diseases. Am J Respir Crit Care Med 175:367–416.

4. Jarand J, Levin A, Zhang L, Huitt G, Mitchell JD, Daley CL. 2011. Clinical and microbiologic outcomes in patients receiving treatment for Mycobacterium abscessus pulmonary disease. Clin Infect Dis 52:565–71.

5. Maurer FP, Bruderer VL, Ritter C, Castelberg C, Bloemberg GV, Bottger EC. 2014. Lack of Antimicrobial Bactericidal Activity in Mycobacterium abscessus. Antimicrob Agents Chemother 58:3828–36.

6. van Ingen J, Boeree MJ, van Soolingen D, Mouton JW. 2012. Resistance mechanisms and drug susceptibility testing of nontuberculous mycobacteria. Drug Resist Updat 15:149–61.

7. Winthrop KL, Marras TK, Adjemian J, Zhang H, Wang P, Zhang Q. 2020. Incidence and Prevalence of Nontuberculous Mycobacterial Lung Disease in a Large U.S. Managed Care Health Plan, 2008-2015. Ann Am Thorac Soc 17:178–185.

8. Dal Molin M, Gut M, Rominski A, Haldimann K, Becker K, Sander P. 2018. Molecular Mechanisms of Intrinsic Streptomycin Resistance in Mycobacterium abscessus. Antimicrob Agents Chemother 62.

9. Hurst-Hess K, Rudra P, Ghosh P. 2017. Mycobacterium abscessus WhiB7 Regulates a Species-Specific Repertoire of Genes To Confer Extreme Antibiotic Resistance. Antimicrob Agents Chemother 61.

10. Nash KA, Brown-Elliott BA, Wallace RJ, Jr. 2009. A novel gene, erm(41), confers inducible macrolide resistance to clinical isolates of Mycobacterium abscessus but is absent from Mycobacterium chelonae. Antimicrob Agents Chemother 53:1367–76.

11. Rudra P, Hurst-Hess K, Lappierre P, Ghosh P. 2018. High Levels of Intrinsic Tetracycline Resistance in Mycobacterium abscessus Are Conferred by a Tetracycline-Modifying Monooxygenase. Antimicrob Agents Chemother 62.

12. Cullen AR, Cannon CL, Mark EJ, Colin AA. 2000. Mycobacterium abscessus infection in cystic fibrosis. Colonization or infection? Am J Respir Crit Care Med 161:641–5.

13. Griffith DE, Aksamit T, Brown-Elliott BA, Catanzaro A, Daley C, Gordin F, Holland SM, Horsburgh R, Huitt G, Iademarco MF, Iseman M, Olivier K, Ruoss S, von Reyn CF, Wallace RJ, Jr., Winthrop K, Subcommittee ATSMD, American Thoracic S, Infectious Disease Society of A. 2007. An official ATS/IDSA statement: diagnosis, treatment, and prevention of nontuberculous mycobacterial diseases. Am J Respir Crit Care Med 175:367–416.

14. Qi LS, Larson MH, Gilbert LA, Doudna JA, Weissman JS, Arkin AP, Lim WA. 2013. Repurposing CRISPR as an RNA-guided platform for sequence-specific control of gene expression. Cell 152:1173–83.

15. Larson MH, Gilbert LA, Wang X, Lim WA, Weissman JS, Qi LS. 2013. CRISPR interference (CRISPRi) for sequence-specific control of gene expression. Nat Protoc 8:2180–96.

16. Akusobi C, Benghomari BS, Zhu J, Wolf ID, Singhvi S, Dulberger CL, Ioerger TR, Rubin EJ. 2022. Transposon mutagenesis in Mycobacterium abscessus identifies an essential penicillin-binding protein involved in septal peptidoglycan synthesis and antibiotic sensitivity. Elife 11.

17. Gupta R, Rohde KH. 2023. Implementation of a mycobacterial CRISPRi platform in Mycobacterium abscessus and demonstration of the essentiality of ftsZ(Mab). Tuberculosis (Edinb) 138:102292.

18. Medjahed H, Reyrat JM. 2009. Construction of Mycobacterium abscessus defined glycopeptidolipid mutants: comparison of genetic tools. Appl Environ Microbiol 75:1331–8.

19. Nguyen TQ, Heo BE, Park Y, Jeon S, Choudhary A, Moon C, Jang J. 2023. CRISPR Interference-Based Inhibition of MAB_0055c Expression Alters Drug Sensitivity in Mycobacterium abscessus. Microbiol Spectr 11:e0063123.

20. Kurepina N, Chen L, Composto K, Rifat D, Nuermberger EL, Kreiswirth BN. 2022. CRISPR Inhibition of Essential Peptidoglycan Biosynthesis Genes in Mycobacterium abscessus and Its Impact on beta-Lactam Susceptibility. Antimicrob Agents Chemother 66:e0009322.

21. Bendixen L, Jensen TI, Bak RO. 2023. CRISPR-Cas-mediated transcriptional modulation: The therapeutic promises of CRISPRa and CRISPRi. Mol Ther 31:1920–1937.

22. Enright AL, Heelan WJ, Ward RD, Peters JM. 2024. CRISPRi functional genomics in bacteria and its application to medical and industrial research. Microbiol Mol Biol Rev 88:e0017022.

23. Fielden J, Siegner SM, Gallagher DN, Schroder MS, Dello Stritto MR, Lam S, Kobel L, Schlapansky MF, Jackson SP, Cejka P, Jost M, Corn JE. 2025. Comprehensive interrogation of synthetic lethality in the DNA damage response. Nature 640:1093–1102.

24. Jana B, Liu X, Denereaz J, Park H, Leshchiner D, Liu B, Gallay C, Zhu J, Veening JW, van Opijnen T. 2024. CRISPRi-TnSeq maps genome-wide interactions between essential and non-essential genes in bacteria. Nat Microbiol 9:2395–2409.

25. Rong Y, Frey A, Ozdemir E, Sainz de la Maza Larrea A, Li S, Nielsen AT, Jensen SI. 2024. CRISPRi-mediated metabolic switch enables concurrent aerobic and synthetic anaerobic fermentations in engineered consortium. Nat Commun 15:8985.

26. Shin J, Bae J, Lee H, Kang S, Jin S, Song Y, Cho S, Cho BK. 2023. Genome-wide CRISPRi screen identifies enhanced autolithotrophic phenotypes in acetogenic bacterium Eubacterium limosum. Proc Natl Acad Sci U S A 120:e2216244120.

27. Sun L, Zheng P, Sun J, Wendisch VF, Wang Y. 2023. Genome-scale CRISPRi screening: A powerful tool in engineering microbiology. Eng Microbiol 3:100089.

28. Zhang Y, Zhang T, Xiao X, Wang Y, Kawalek A, Ou J, Ren A, Sun W, de Bakker V, Liu Y, Li Y, Yang L, Ye L, Jia N, Veening JW, Liu X. 2025. CRISPRi screen identifies FprB as a synergistic target for gallium therapy in Pseudomonas aeruginosa. Nat Commun 16:5870.

29. Bosch B, DeJesus MA, Poulton NC, Zhang W, Engelhart CA, Zaveri A, Lavalette S, Ruecker N, Trujillo C, Wallach JB, Li S, Ehrt S, Chait BT, Schnappinger D, Rock JM. 2021. Genome-wide gene expression tuning reveals diverse vulnerabilities of M. tuberculosis. Cell 184:4579–4592 e24.

30. Blumenthal A, Trujillo C, Ehrt S, Schnappinger D. 2010. Simultaneous analysis of multiple Mycobacterium tuberculosis knockdown mutants in vitro and in vivo. PLoS One 5:e15667.

31. Boldrin F, Ventura M, Degiacomi G, Ravishankar S, Sala C, Svetlikova Z, Ambady A, Dhar N, Kordulakova J, Zhang M, Serafini A, Vishwas KG, Kolly GS, Kumar N, Palu G, Guerin ME, Mikusova K, Cole ST, Manganelli R. 2014. The phosphatidyl-myo-inositol mannosyltransferase PimA is essential for Mycobacterium tuberculosis growth in vitro and in vivo. J Bacteriol 196:3441–51.

32. Gandotra S, Schnappinger D, Monteleone M, Hillen W, Ehrt S. 2007. In vivo gene silencing identifies the Mycobacterium tuberculosis proteasome as essential for the bacteria to persist in mice. Nat Med 13:1515–20.

33. Leblanc C, Prudhomme T, Tabouret G, Ray A, Burbaud S, Cabantous S, Mourey L, Guilhot C, Chalut C. 2012. 4’-Phosphopantetheinyl transferase PptT, a new drug target required for Mycobacterium tuberculosis growth and persistence in vivo. PLoS Pathog 8:e1003097.

34. Marrero J, Rhee KY, Schnappinger D, Pethe K, Ehrt S. 2010. Gluconeogenic carbon flow of tricarboxylic acid cycle intermediates is critical for Mycobacterium tuberculosis to establish and maintain infection. Proc Natl Acad Sci U S A 107:9819–24.

35. Puckett S, Trujillo C, Eoh H, Marrero J, Spencer J, Jackson M, Schnappinger D, Rhee K, Ehrt S. 2014. Inactivation of fructose-1,6-bisphosphate aldolase prevents optimal co-catabolism of glycolytic and gluconeogenic carbon substrates in Mycobacterium tuberculosis. PLoS Pathog 10:e1004144.

36. Weiss LA, Stallings CL. 2013. Essential roles for Mycobacterium tuberculosis Rel beyond the production of (p)ppGpp. J Bacteriol 195:5629–38.

37. Woong Park S, Klotzsche M, Wilson DJ, Boshoff HI, Eoh H, Manjunatha U, Blumenthal A, Rhee K, Barry CE, 3rd, Aldrich CC, Ehrt S, Schnappinger D. 2011. Evaluating the sensitivity of Mycobacterium tuberculosis to biotin deprivation using regulated gene expression. PLoS Pathog 7:e1002264.

38. Bernut A, Herrmann JL, Ordway D, Kremer L. 2017. The Diverse Cellular and Animal Models to Decipher the Physiopathological Traits of Mycobacterium abscessus Infection. Front Cell Infect Microbiol 7:100.

39. Bernut A, Le Moigne V, Lesne T, Lutfalla G, Herrmann JL, Kremer L. 2014. In vivo assessment of drug efficacy against Mycobacterium abscessus using the embryonic zebrafish test system. Antimicrob Agents Chemother 58:4054–63.

40. Boudehen YM, Tasrini Y, Aguilera-Correa JJ, Alcaraz M, Kremer L. 2023. Silencing essential gene expression in Mycobacterium abscessus during infection. Microbiol Spectr 11:e0283623.

41. Luo D, Xie W, Wang C, Sun Y, Zhang L, Qian L, Zhang J, Dang G, Liu S, Wang Z. 2025. Isoleucyl-tRNA synthetase depletion reveals vulnerabilities in Mycobacterium abscessus and Mycobacterium marinum. Commun Biol 8:1379.

42. Xie W, Luo D, Wu M, Sun Y, Wang Z. 2025. The evaluation of Phenylalanine-tRNA ligase beta unit (PheT), as a potential target in Mycobacterium abscessus. Tuberculosis (Edinb) 152:102626.

43. Rock JM, Hopkins FF, Chavez A, Diallo M, Chase MR, Gerrick ER, Pritchard JR, Church GM, Rubin EJ, Sassetti CM, Schnappinger D, Fortune SM. 2017. Programmable transcriptional repression in mycobacteria using an orthogonal CRISPR interference platform. Nat Microbiol 2:16274.

44. DeJesus MA, Gerrick ER, Xu W, Park SW, Long JE, Boutte CC, Rubin EJ, Schnappinger D, Ehrt S, Fortune SM, Sassetti CM, Ioerger TR. 2017. Comprehensive Essentiality Analysis of the Mycobacterium tuberculosis Genome via Saturating Transposon Mutagenesis. mBio 8.

45. Rifat D, Chen L, Kreiswirth BN, Nuermberger EL. 2021. Genome-Wide Essentiality Analysis of Mycobacterium abscessus by Saturated Transposon Mutagenesis and Deep Sequencing. mBio 12:e0104921.

46. Rimal B, Lippincott CK, Panthi CM, Xie Y, Keepers TR, Alley M, Lamichhane G. 2024. Efficacy of epetraborole against Mycobacteroides abscessus in a mouse model of lung infection. Antimicrob Agents Chemother 68:e0064824.

47. Singh V, Dziwornu GA, Mabhula A, Chibale K. 2021. Rv0684/fusA1, an Essential Gene, Is the Target of Fusidic Acid and Its Derivatives in Mycobacterium tuberculosis. ACS Infect Dis 7:2437–2444.

48. Sullivan JR, Lupien A, Kalthoff E, Hamela C, Taylor L, Munro KA, Schmeing TM, Kremer L, Behr MA. 2021. Efficacy of epetraborole against Mycobacterium abscessus is increased with norvaline. PLoS Pathog 17:e1009965.

49. Wu W, He S, Li A, Guo Q, Tan Z, Liu S, Wang X, Zhang Z, Li B, Chu H. 2022. A Novel Leucyl-tRNA Synthetase Inhibitor, MRX-6038, Expresses Anti-Mycobacterium abscessus Activity In Vitro and In Vivo. Antimicrob Agents Chemother 66:e0060122.

50. Miller MA, Stabenow JM, Parvathareddy J, Wodowski AJ, Fabrizio TP, Bina XR, Zalduondo L, Bina JE. 2012. Visualization of murine intranasal dosing efficiency using luminescent Francisella tularensis: effect of instillation volume and form of anesthesia. PLoS One 7:e31359.

51. Southam DS, Dolovich M, O’Byrne PM, Inman MD. 2002. Distribution of intranasal instillations in mice: effects of volume, time, body position, and anesthesia. Am J Physiol Lung Cell Mol Physiol 282:L833–9.

52. Ebino K, Lemus R, Karol MH. 1999. The importance of the diluent for airway transport of toluene diisocyanate following intranasal dosing of mice. Inhal Toxicol 11:171–85.

53. Gengenbacher M, Zimmerman MD, Sarathy JP, Kaya F, Wang H, Mina M, Carter C, Hossen MA, Su H, Trujillo C, Ehrt S, Schnappinger D, Dartois V. 2020. Tissue Distribution of Doxycycline in Animal Models of Tuberculosis. Antimicrob Agents Chemother 64.

54. Lim B, Zimmermann M, Barry NA, Goodman AL. 2017. Engineered Regulatory Systems Modulate Gene Expression of Human Commensals in the Gut. Cell 169:547–558 e15.

55. Bosch B, DeJesus MA, Poulton NC, Zhang W, Engelhart CA, Zaveri A, Lavalette S, Ruecker N, Trujillo C, Wallach JB, Li S, Ehrt S, Chait BT, Schnappinger D, Rock JM. 2021. Genome-wide gene expression tuning reveals diverse vulnerabilities of M. tuberculosis. Cell doi:10.1016/j.cell.2021.06.033.

56. Gibson AJ, Passmore IJ, Faulkner V, Xia D, Nobeli I, Stiens J, Willcocks S, Clark TG, Sobkowiak B, Werling D, Villarreal-Ramos B, Wren BW, Kendall SL. 2021. Probing Differences in Gene Essentiality Between the Human and Animal Adapted Lineages of the Mycobacterium tuberculosis Complex Using TnSeq. Front Vet Sci 8:760717.

57. Rohde KH, Abramovitch RB, Russell DG. 2007. Mycobacterium tuberculosis invasion of macrophages: linking bacterial gene expression to environmental cues. Cell Host Microbe 2:352–64.

